# The HCN1 hyperpolarization-activated cyclic nucleotide-gated channel enhances evoked GABA release from parvalbumin positive interneurons

**DOI:** 10.1101/2022.11.11.516205

**Authors:** Tobias Bock, Eric W. Buss, Olivia M. Lofaro, Felix Leroy, Bina Santoro, Steven A. Siegelbaum

## Abstract

Hyperpolarization-activated, cyclic nucleotide-gated (HCN) channels generate the cationic Ih current in neurons and regulate the excitability of neuronal networks. The function of HCN channels depends, in part, on their subcellular localization. Of the four HCN isoforms (HCN1-4), HCN1 is strongly expressed in the dendrites of pyramidal neurons in hippocampal area CA1 but also in presynaptic terminals of parvalbumin-positive interneurons (PV+ INs), which provide strong inhibitory control over hippocampal activity. Yet, little is known about how HCN1 channels in these cells regulate the evoked release of the inhibitory transmitter GABA from their axon terminals. Here, we used several genetic, optogenetic, electrophysiological and imaging techniques to investigate how the electrophysiological properties of PV+ INs are regulated by HCN1, including how HCN1 activity at presynaptic terminals regulates the release of GABA onto pyramidal neurons (PNs) in CA1. We found that application of HCN1 pharmacological blockers reduced the amplitude of the inhibitory postsynaptic potential recorded from CA1 pyramidal neurons in response to selective optogenetic stimulation of PV+ INs. Homozygous HCN1^-/-^ knockout mice also show reduced IPSCs in postsynaptic cells. Finally, two-photon imaging using genetically encoded fluorescent calcium indicators revealed that HCN1 blockers reduced the probability that an extracellular electrical stimulating pulse evoked a Ca^2+^ response in individual PV+ IN presynaptic boutons. Taken together, our results show that HCN1 channels in the axon terminals of PV+ interneurons facilitate GABAergic transmission in the hippocampal CA1 region.

## Introduction

A wide range of GABAergic inhibitory interneurons contribute to shaping neuronal network activity throughout the brain. Their axons commonly target precise locations in various subcellular compartments of their postsynaptic cells, defining their specific role in modulating excitatory activity (Klausberger and Somogyi, 2008; Pelkey et al., 2017). In the hippocampus and neocortex, somatostatin expressing interneurons target distal dendritic regions of excitatory pyramidal neurons (PNs), where they regulate synaptic integration and plasticity (Chiu et al., 2013; Maccaferri, 2005; Schulz et al., 2018; Sun et al., 2014). In contrast, parvalbumin expressing interneurons (PV+ INs) target the perisomatic subdomain and axon initial segment of PNs, thereby regulating spike output and network oscillation (Geiger et al., 1997; Pouille and Scanziani, 2001; Sohal et al., 2009; Sun et al., 2014).

While the synaptic connections that interneurons make with their targets define the network activity that ultimately determines behavioral output (Freund, 2003; Lapray et al., 2012), the efficacy of these inhibitory connections depends on the makeup of their voltage-gated ion channels in distinct subcellular compartments, including soma, dendrites, axons and presynaptic boutons. The clinical importance of many of these voltage-gated channels is indicated by findings that, in humans and animal models, mutations in interneuron channels can lead to different forms of epileptic encephalopathies (Katsarou et al., 2017). Perhaps the best understood example is Dravet’s syndrome, where a loss-of-function mutation in the SCN1A excitatory voltage-gated sodium channel, which is strongly expressed in PV+ INs, causes seizure activity due to a reduction in PV+ IN excitability, resulting in decreased inhibition (Kaneko et al., 2022; Tai et al., 2014). Mutations in the HCN1 subtype of the hyperpolarization-activated, cyclic nucleotide-modulated (HCN) channel family, which is strongly expressed in PV+ IN axons and presynaptic boutons, have recently been found to underlie certain cases of early infantile epileptic encephalopathy (EIEE; Kessi et al., Marini et al., 2018; 2022; Nava et al., 2014). However, our understanding of how HCN1 channels contribute to interneuron function remains relatively unexplored.

The family of HCN channels generate the cationic current Ih, which dynamically controls membrane resting potential, input resistance and synaptic integration at post-synaptic sites, thereby regulating the excitability of neuronal networks (Biel et al., 2009; Nolan et al., 2004; Nolan et al., 2003; Sartiani et al., 2017). The channels are encoded by four closely related HCN genes (HCN1-4), which display distinct patterns of expression, with HCN1 expressed in select brain regions, in particular in the neocortex, hippocampus and cerebellum (Lorincz et al., 2002; Nolan et al., 2003; Rinaldi et al., 2013; Santoro and Shah, 2020).

In the hippocampus HCN1 is strongly expressed in CA1 pyramidal neurons, where it is localized to apical dendrites forming a gradient of increasing density with increasing distance from the soma (Lorincz et al., 2002; Magee, 1998; Notomi and Shigemoto, 2004). There HCN1 acts as an inhibitory constraint on the temporal integration of the perforant path (PP) excitatory postsynaptic potentials (EPSPs) CA1 PNs receive from entorhinal cortex, which terminate on the CA1 distal apical dendrites, the site of highest HCN1 expression (Magee, 1999). HCN1 also suppresses long-term potentiation of the PP EPSP and constrains hippocampal-dependent spatial reference memory (Nolan et al., 2004). At the systems level of hippocampal function, loss of HCN1 alters the spatial coding properties of both CA1 and CA3 PNs, resulting in a decreased precision of spatial firing (increased place field size) but enhanced day-to-day stability of place field location (Hussaini et al., 2011). In contrast, specific deletion of HCN1 in entorhinal cortex decreases grid cell spatial stability, associated with impaired learning and memory.

In addition to its role in regulating pyramidal neuron function, HCN1 is also strongly expressed in parvalbumin-positive inhibitory neurons (PV+ INs), including hippocampal basket cells (Notomi and Shigemoto, 2004; Santoro et al., 1997). Thus, to understand the role HCN channels at a systems level, it is important to study their role in regulating PV+ IN activity, as this class of neurons represent ~24% of GABAergic interneurons in the hippocampus (Bezaire and Soltesz, 2013; Deng et al.). Moreover, PV+ INs have been implicated in generating hippocampal network activity, including gamma and theta oscillations, and have been implicated in controlling hippocampal spatial coding, object and spatial recognition memory as well as social memory (Caroni, 2015; Deng et al., 2019; Fuchs et al., 2007; Korotkova et al., 2010; Murray et al., 2011; Wulff et al., 2009). Furthermore, PV+ IN function is of increasing clinical relevance. Due to their high firing frequency, PV+ INs are particularly vulnerable to metabolic and oxidative stress, and impairments in the function or connectivity of PV+ INs have been associated with epilepsy, schizophrenia, autism and cognitive decline (Benes, 2015; Jiang et al., 2016; Kann, 2016).

The role of HCN1 is particularly intriguing in PV+ INs, where the channel is found predominantly in the axon and presynaptic terminals. Axonal HCN channels are indeed critical regulators of PV+ IN excitability, as they have been shown to enhance action potential (AP) initiation during sustained high-frequency firing and facilitate the propagation of AP trains (Roth and Hu, 2020), in addition to being required for the maintenance of persistent firing (Elgueta et al., 2015). Prior studies have also suggested that Ih and HCN1 in PV+ INs may act to promote both miniature (Aponte et al., 2006) and spontaneous (Lupica et al., 2001) inhibitory postsynaptic potentials (IPSPs), based on the finding that pharmacological blockers of Ih can reduce the rate of miniature and spontaneous IPSPs. However, these studies did not examine the contributions of HCN channels to evoked IPSPs. Nor did they distinguish potential effects of HCN channel blockers on IPSPs from non-PV+ INs or the contribution of HCN1 from other HCN isoforms. Importantly, these blockers can have significant off-target effects that can alter release independently of HCN channels (Chevaleyre and Castillo, 2002). Thus, the role of HCN1 in PV+ IN function remains uncertain. Here we used a combination of pharmacological, genetic and calcium imaging-based approaches to identify the role of HCN1 both in the regulation of PV+ IN somatic properties and in the control of the efficacy of inhibitory synaptic transmission from PV+ INs onto their CA1 PN targets.

## Results

### HCN1 is expressed in PV+ IN presynaptic boutons

To determine which subunit is most relevant for HCN channel formation in presynaptic terminals of PV+ interneurons within CA1, we utilized immunofluorescent labeling. We colabeled hippocampal slices with antibodies against synaptotagmin-2 (SYT2), a calcium sensor, localized on synaptic vesicles of PV+ INs (Garcia-Junco-Clemente et al., 2010; Sommeijer and Levelt, 2012) and antibodies against either the HCN1 or HCN2 subunits. Both, HCN1 and HCN2 labeling showed expression in CA1, mainly in *stratum radiatum* (SR) and *stratum lacunosum moleculare* (SLM), where HCN channels are found at high density within the apical dendrites of CA1 pyramidal neurons (Figure 1). However, HCN1 labeling also revealed expression within *stratum pyramidale* (SP), where it showed a more diffuse perisomatic expression pattern that co-localized with SYT2 staining (Figure 1a-c). In contrast, immunostaining for HCN2 subunits was weak in SP and not co-localized with SYT2, indicating that it is primarily HCN1 subunits, which form the HCN channels present in the presynaptic terminals in SP (Figure 1d-f).

**Figure 1.**
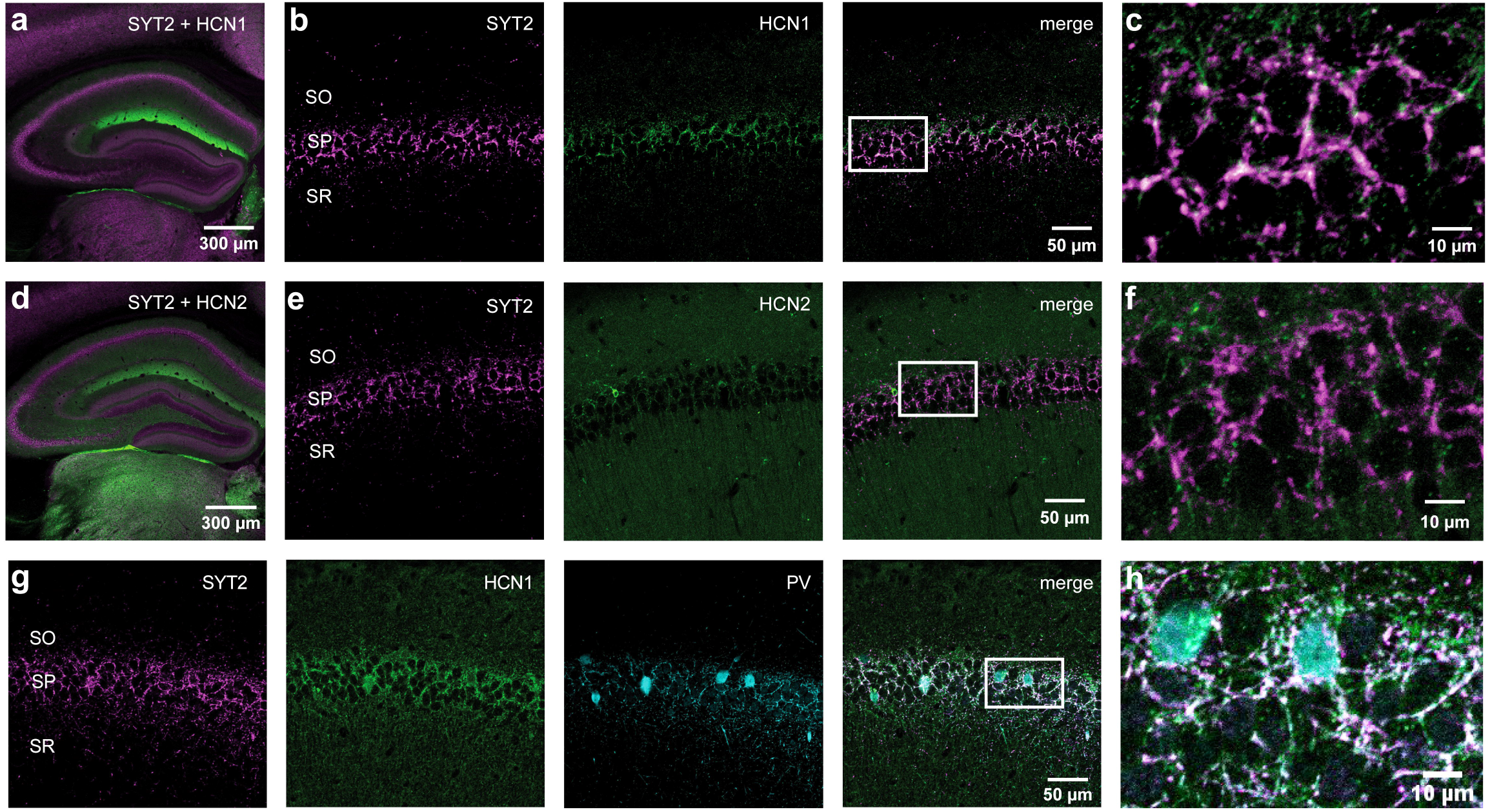
Confocal images of double and triple immunofluorescence staining in hippocampus. **(a)** Low magnification overlay image of dual staining for synaptotagmin 2 (SYT 2, purple) and HCN1 (green) of the entire hippocampus **(b)** High-magnification images of area CA1, showing *stratum oriens* (SO), *stratum pyramidale* (SP) and *stratum radiatum* (SR); left: stained for synaptotagmin 2 (SYT2, purple); middle: stained for HCN1 (green); right: overlay of the two stainings (colocalization shown in white). **(c)** Close-up of the white square outline of the overlay image in **(b). (d)** Low magnification overlay image of dual staining for synaptotagmin 2 (SYT 2, purple) and HCN2 (green) of the entire hippocampus **(e)** High-magnification images of area CA1, showing *stratum oriens* (SO), *stratum pyramidale* (SP) and *stratum radiatum* (SR); left: stained for synaptotagmin 2 (SYT2, purple); middle: stained for HCN2 (green); right: overlay of the two stainings (colocalization shown in white). **(f**) Close-up of the white square outline of the overlay image in **(e). (g)** High-magnification images of area CA1, showing *stratum oriens* (SO), *stratum pyramidale* (SP) and *stratum radiatum* (SR); far left: stained for synaptotagmin 2 (SYT2, purple); middle left: stained for HCN1 (green); middle right: stained for parvalbumin (PV, cyan); far right: overlay of the three stainings (colocalization shown in white). **(h)** Close-up of the white square outline of the overlay image in **(g).**

While SYT2 has been shown to be expressed specifically in GABAergic interneurons both in the cortex (Sommeijer and Levelt, 2012) and in hippocampal cell cultures (Garcia-Junco-Clemente et al., 2010), we also performed triple labeling for PV, SYT2 and HCN1 to confirm the localization of HCN1 to PV+ INs. Indeed we found that all three markers were colocalized in SP within axonal boutons. In contrast, PV+ IN somas, defined by PV antibody staining, lacked any detectable staining for either HCN1 or SYT2 (Figure 1g-h). Thus we conclude that HCN1 is indeed is localized in the axons and presynaptic boutons of PV+ INs within the CA1 SP layer.

### HCN channels have little impact on PV+ IN somatic membrane properties

To determine the impact of HCN channels on cellular excitability and membrane properties in PV+ INs, we crossed a mouse line expressing Cre recombinase selectively in PV+ INs (*Pvalb^tm1(cre)Arbr^/J*, PV-Cre hereafter) with a reporter mouse line, which expresses the fluorescent marker tdTomato in a Cre-dependent manner (*GT(ROSA)^26Sortm14(CAG-tdTomato)Hze^/J*, or Ai14). This resulted in reliable tdTomato expression in PV+ INs specifically (Figure 2a,b), which allowed us to perform whole-cell patch-clamp recordings from identified PV+ INs in the CA1 region of acute hippocampal slices. These experiments revealed two populations of PV+ INs based on their electrophysiological properties using whole cell current clamp conditions.

**Figure 2.**
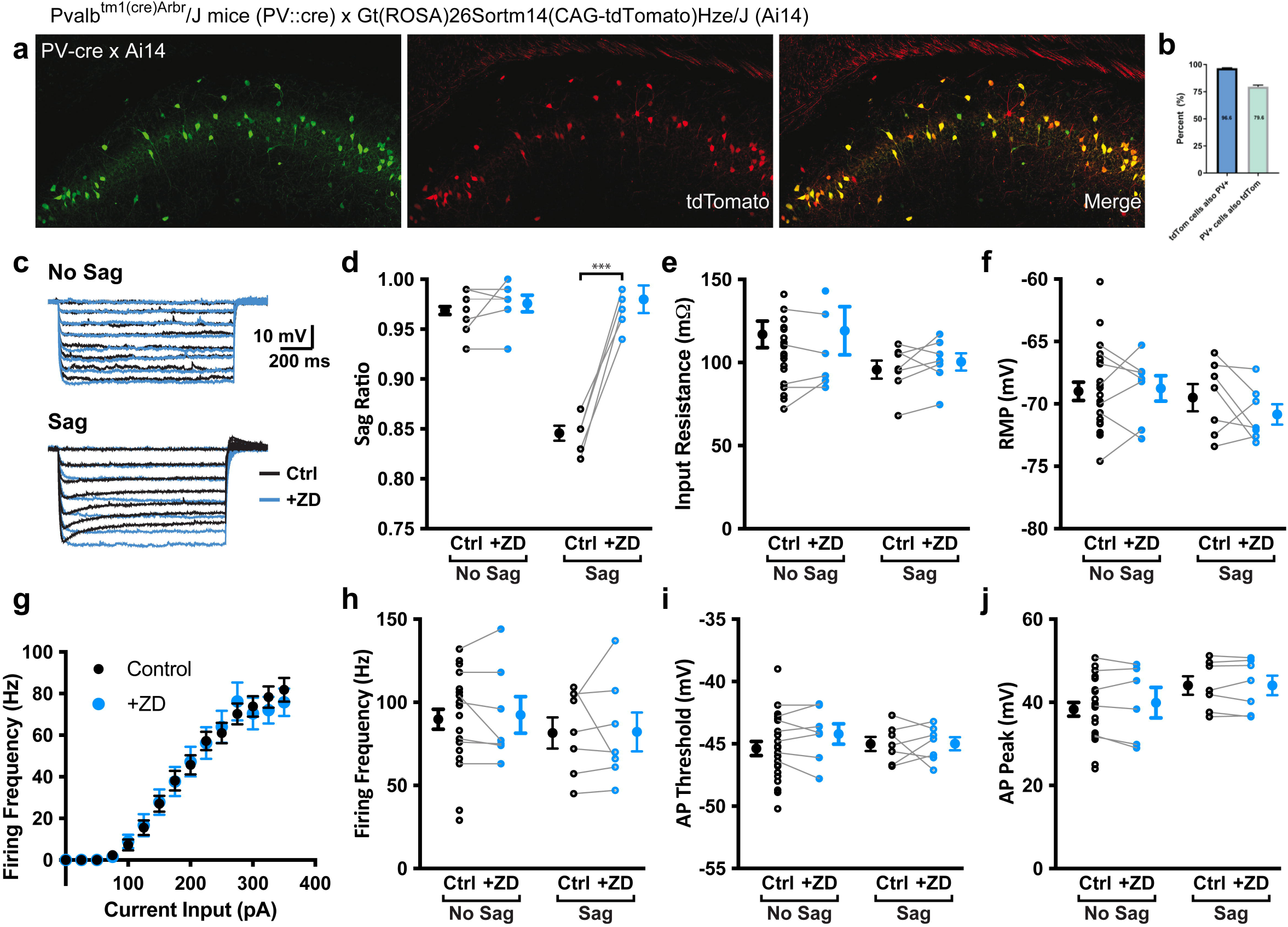
Impact of HCN channel block on somatic properties of PV+ INs. **(a)** Confocal section of parvalbumin immunofluorescence staining and tdTomato expression levels after Cre recombination. **(b)** quantification of colocalization of tdtTomato and PV. **(c)** Example traces of current steps from 0 to −175 pA (in steps of −25 pA) of a cell exhibiting no voltage sag (top) and one with sag (bottom) before (black) and after (blue) bath application of ZD7288. **(d-f)** Summary graphs of the effects of ZD 7288 (before: black, after: blue) on passive membrane parameters, divided by “sag” and “non-sag” cells. **(g)** AP firing frequency in response to positive current steps (F-I curve) before (black) and after (blue) ZD7288. **(h-j)** Summary graphs of the effects of ZD 7288 (before: black, after: blue) on action potential properties, divided by sag and non-sag cells.

The majority of PV+ cells from which we obtained whole-cell recordings (22 out of 29) showed no voltage sag - a hallmark of HCN channel activation - in response to hyperpolarizing current steps (“no sag” cells). The remainder of PV+ INs (7 out of 29) exhibited a moderate voltage sag (“sag” cells, Figure 2c). We quantified the extent of sag by measuring the ratio of the membrane voltage at the peak of the hyperpolarization at the start of the current pulse to the steady-state membrane voltage at the end of the 1-s long current pulse (sag ratio).

Bath application of the HCN blocker ZD7288 (10 μM) blocked the sag in the sag cell population (sag ratio = 0.85 ± 0.01 before compared to 0.98 ± 0.01 after ZD7288; p < 0.001 with paired t-test; n = 7; Figure 2d). ZD7288 had no effect on the sag ratio of the non-sag cells (0.97 ± 0.01 before vs. 0.98 ± 0.01 after ZD7288; p = 0.37; n = 22).

Somewhat unexpectedly, ZD7288 had no effect in sag cells on input resistance (95.71 ± 5.4 MΩ before compared to 100.39 ± 5.2 MΩ after ZD7288; p = 0.30; n = 7; Figure 2e) or resting membrane potential (−69.5 ± 1.1 mV before compared to −70.8 ± 0.8 mV after ZD7288; p = 0.22; n = 7; Figure 2f), even though HCN channels help lower input resistance and depolarize the resting membrane in many cell types in which they are expressed (Robinson and Siegelbaum, 2003). This suggests that the HCN channels may be only weakly expressed in the soma of PV+ INs, even in sag cells.

Bath application of ZD7288 had no impact on action potential (AP) properties, such as firing frequency (81.57 ± 9.4 Hz before compared to 82.14 ± 11.69 Hz after ZD7288 with a +350 pA current step; p = 0.57; n = 7), threshold (−45.0 ± 5.7 mV before compared to −45.0 ± 5.6 Hz after ZD7288; p = 0.98; n = 7) or peak voltage (44.1 ± 2.2 mV before compared to 44.1 ± 2.3 mV after ZD7288; p = 0.98; n = 7). From these data, we conclude that, although there may be two distinct populations of PV+ INs in CA1, HCN channels do not have a significant influence on resting membrane properties, apart from generating a moderate sag in a subset of cells, or on somatic action potential firing for either PV+ IN population.

### HCN channels regulate the evoked inhibitory postsynaptic current in CA1 pyramidal neurons

The fact that we saw strong HCN1 expression in the synaptic terminals of PV+ INs in CA1 SP (Figure 1) suggests that the main impact of HCN channels in this cell type may be on synaptic transmission, rather than on somatic membrane properties. To explore this possibility, we measured the effect that blockade of HCN channels with ZD7288 exerted on the inhibitory postsynaptic currents (IPSCs) evoked by electrical stimulation in the CA1 SP layer and recorded in CA1 PNs in voltage-clamp configuration. To activate directly local INs we placed the stimulating electrode in SP relatively close to the recorded CA1 PN (150 to 300 μm from the recorded PN). To examine only monosynaptic IPSCs we blocked fast excitatory synaptic transmission by applying the AMPA receptor antagonist CNQX and the NMDA receptor blocker APV to the bath solution. To focus on effects of HCN block on presynaptic INs we included the intracellular HCN channel blocker QX-314 in the internal solution of the patch pipette to block the effects of postsynaptic HCN channels in the CA1 PN membrane. IPSCs were recorded while the cell was clamped to +10 mV before and after bath application of ZD7288 (Figure 3a). We applied ZD7288 at 10 mM and limited the duration of its application to 10 min to minimize off-target effects (Chevaleyre and Castillo, 2002).

**Figure 3.**
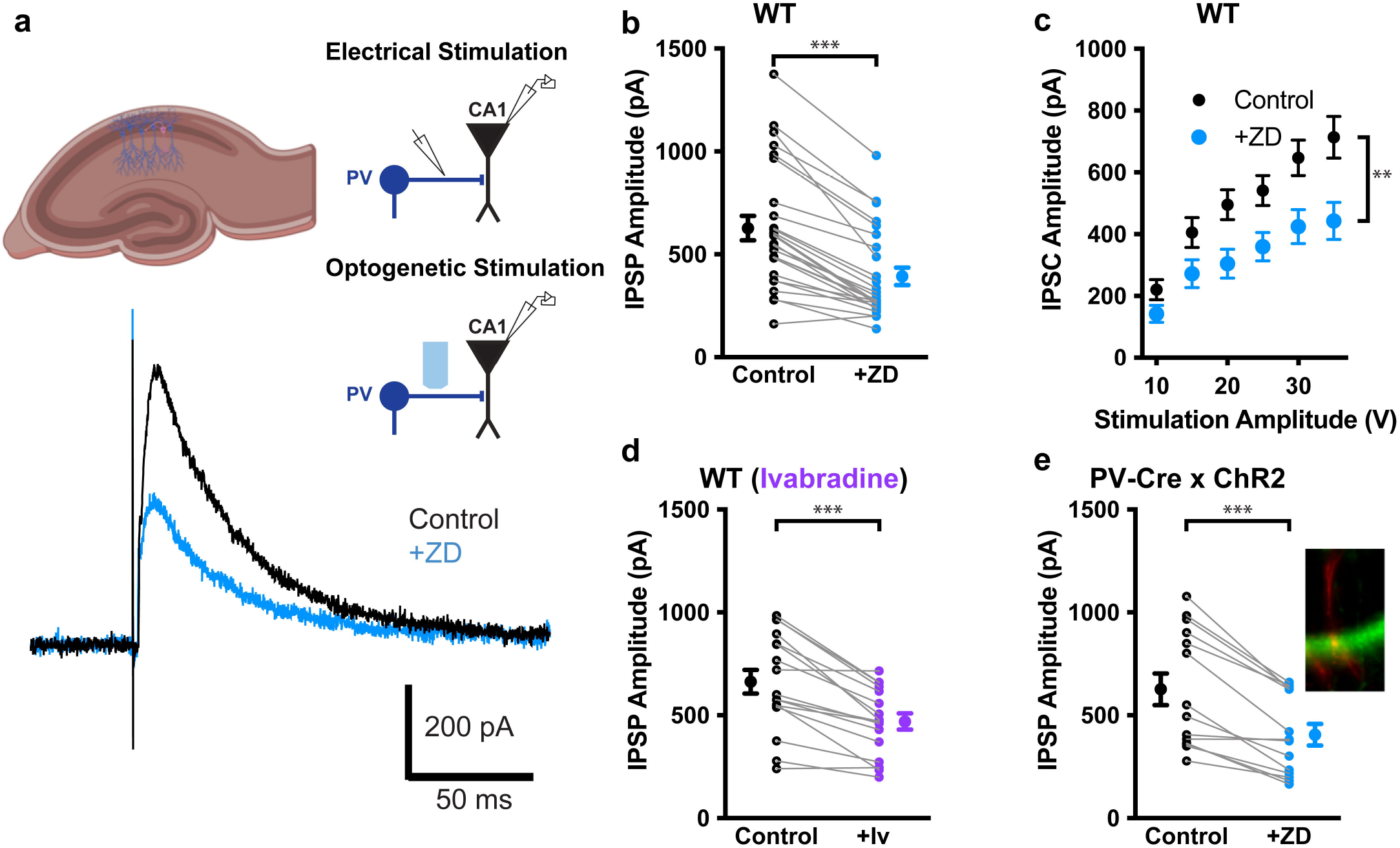
Block of HCN channels decreases IPSCs evoked by stimulation of PV+ INs. **(a)** (top) Schematic of voltage clamp recordings from CA1 pyramidal neurons with a stimulating electrode in the pyramidal cell layer or optogenetically stimulating using blue light pulses (2ms). (bottom) Example trace of an IPSC before (back) and after (blue) ZD7288 (10 μM) application. Excitatory transmission was blocked with CNQX (25 μM) and APV (50 μM). QX-314 was used in the recording pipette to block postsynaptic HCN channels. **(b)** Evoked IPSCs using extracellular electrical stimulation before (black) and after (blue) bath application of ZD7288. (c) Input-output curve of IPSC amplitude as function of voltage stimulus intensity, ranging from 10-35V before (black) and after (blue) bath application of ZD7288. **(d)** Evoked IPSCs using extracellular electrical stimulation before (black) and after (purple) bath application of ivabradine (30 μM). **(e)** Light pulse stimulation of PV+ IN axons expressing ChR2 before (black) and after (blue) bath application of ZD7288. Insert shows a pyramidal neuron filled with biocytin in red and ChR2, expressed in PV INs in green.

We found that application of the HCN channel blocker ZD7288 caused a significant ~35% reduction in IPSC amplitude (from 627.3 ± 58.0 pA before to 393.1 ± 41.4 pA after ZD7288 at a stimulation amplitude of 35 V; p < 0.0001; n = 26; Figure 3b,c). In addition, the more selective HCN channel blocker ivabradine exerted a similar inhibitory effect (IPSC reduced from 663.1 ± 57.2 pA to 469.8 ± 39.3 pA after ivabradine application; p < 0.0001; n = 17; Figure 3d), indicating that the reduction of the IPSC after ZD7288 application is not due to off-target effects of the drug.

As extracellular electrical stimulation recruits a variety of IN subtypes, not just PV+ INs, we used an optogenetic approach to selectively activate the PV+ INs. Thus, we crossed the PV-Cre mouse line with a mouse line that expresses channelrhodopsin-2 (ChR2) in a Cre-dependent manner (*Gt(ROSA)26Sor^tm32(CAG-COP4*H134R/EYFP)Hze^/J*, or Ai32), resulting in ChR2 expression specifically in PV+ INs (Figure 3d, insert). We then stimulated these cells and their axons with a 2-ms light pulse at 470 nm, which evoked large IPSCs in CA1 PNs. Similar to its effect on the electrically-evoked IPSC, bath application of ZD-7288 caused a significant decrease in the optogenetically evoked IPSC, from 626 ± 76.8 pA in the absence of drug to 405.8 ± 52.5 pA in the presence of drug (p < 0.0001; n = 14; Figure 3e), confirming that HCN channels expressed in PV+ INs help regulate inhibitory synaptic transmission.

### Primary importance of the HCN1 isoform in regulating inhibitory synaptic transmission

To gain insight into whether it is the HCN1 channel isoform that contributes to the efficacy of inhibitory synaptic transmission mediated by PV+ INs, we examined the effect of ZD7288 on IPSCs recorded in an unconditional HCN1 knockout mouse line (Hcn1^tm2Kndl^/J, Nolan et al. 2003). IPSCs were recorded from CA1 PNs in hippocampal slices from the HCN1^-/-^ mice in response to electrical stimulation in the SP layer, in the presence of CNQX and APV in the extracellular solution.

As expected, application of ZD7288 caused a reduction in IPSC amplitude in wild-type (HCN1^+/+^) littermates (IPSC = 940.5 ± 92.4 pA before and 721.8 ± 92.8 pA after ZD7288; p = 0.0029; n = 8; Figure 4a). The HCN blocker also produced a significant reduction in the IPSC in heterozygous mice missing one HCN1 allele (HCN1^+/-^; IPSC = 912.4 ± 85.0 pA before and 693.6 ± 76.2 pA after ZD7288; p = 0.0001; n = 9; Figure 4b). In contrast, application of ZD7288 caused only a small ~5% decrease in the IPSC in homozygous knockout mice (HCN1^-/-^;IPSC = 626.4 ± 65.8 pA before and 597.9 ± 63.9 pA after ZD7288; p = 0.056; n = 14; Figure 4c). This result demonstrates that the action of the HCN channel blocker to reduce the IPSC was mediated by a specific effect on channels containing the HCN1 subunit.

**Figure 4.**
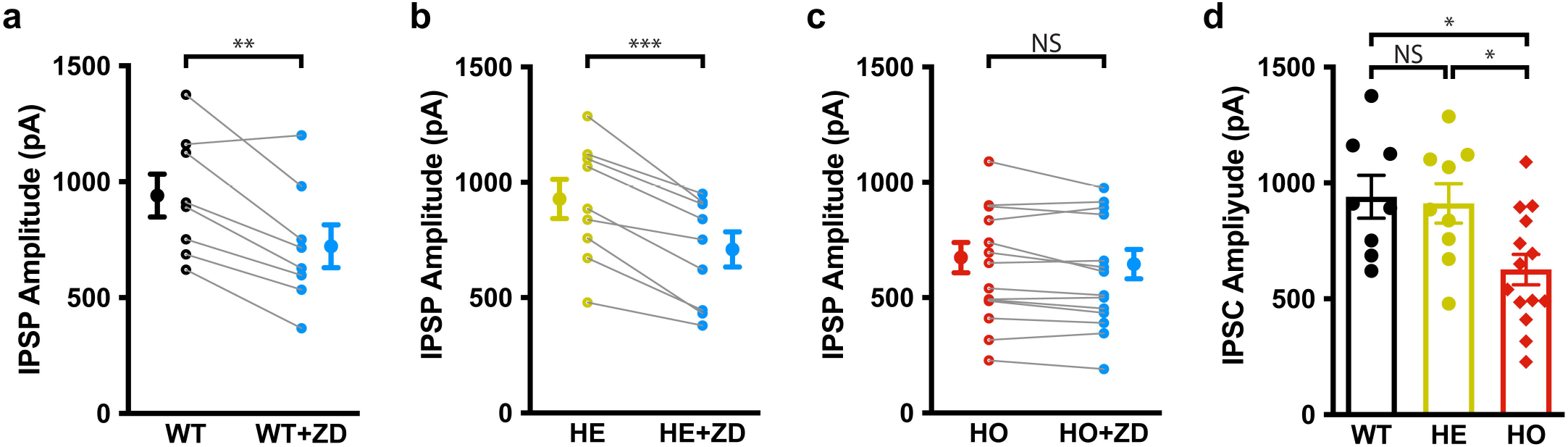
The impact of HCN channel block on IPSCs is occluded in homozygous HCN1 knockout mice. **(a)** Evoked IPSCs using extracellular electrical stimulation before (black) and after (blue) bath application of ZD7288 in wild-type (HCN1^+/+^) littermates of HCN1 KO mice. **(b)** Evoked IPSCs using extracellular electrical stimulation before (yellow) and after (blue) bath application of ZD7288 in heterozygous (HCN1^+/-^) mice. **(c)** Evoked IPSCs using extracellular electrical stimulation before (red) and after (blue) bath application of ZD7288 in homozygous (HCN1^-/-^) mice. **(d)** Comparison of the evoked IPSC amplitudes of wild-type (black), HCN1^+/-^ (yellow) and HCN1^-/-^ (red) mice before bath application ZD7288.

If HCN1 does indeed play an important role in controlling the strength of synaptic inhibition, then homozygous loss of this subunit should lead to a reduction in the amplitude of the IPSC relative to that in wild-type animals. Indeed, we found that the amplitude of the IPSC in HCN1^-/-^ animals (626.4 ± 65.8 pA; n=14) was significantly smaller than that in either wild-type littermates (940.5 ± 92.4 pA; n=8; p = 0.016 with Mann-Whitney test after 1-way ANOVA) or heterozygotes (912.4 ± 85.0 pA; p = 0.028; n=9). In contrast, there was no difference in IPSC size between wild-type and heterozygous mice (p=0.74; Figure 4d). These data indicate that one HCN1 allele enables the cell to express a sufficient level of HCN channels in the presynaptic terminals to maintain a normal-sized IPSC.

### An HCN channel pharmacological antagonist increases the paired-pulse ratio due to blockade of HCN1

How exactly do HCN channels modulate synaptic transmission? To investigate this question further, we measured IPSCs evoked by paired pulse stimulation (50 ms interpulse-interval, Figure 5a). The magnitude of the paired-pulse ratio (PPR) is thought to be inversely related to the probability of transmitter release, as a higher initial release probability during the first pulse will lead to a greater transient depletion of docked vesicles, lowering the probability of vesicle release during the second pulse. We found that application of ZD7288 caused a significant increase in the paired-pulse ratio for IPSCs, evoked by either extracellular electrical stimulation in SP (PPR = 0.28 ± 0.03 in absence and 0.43 ± 0.02 in presence of ZD7288; n=18; p < 0.0001; Figure 5b) or photostimulation of ChR2-expressing PV+ INs (PPR = 0.30 ± 0.03 in absence and 0.47 ± 0.06 in presence of ZD7288; n=8; p = 0.011; Figure 5c). We also observed an increase in PPR with ZD7288 application in heterozygous HCN1 knockout mice (PPR= 0.24 ± 0.02 in absence and 0.30 ± 0.02 in presence of ZD7288; n=9; p = 0.012; Figure 5d). In contrast, ZD7288 had no effect on PPR for IPSCs elicited by electrical stimulation in HCN1^-/-^ mice (PPR = 0.31 ± 0.03 in absence and 0.33 ± 0.03 in presence of ZD7288; p = 0.10; n=14; Figure 5e). These results indicate that presynaptic HCN1 channels in PV+ INs likely enhance the magnitude of the IPSC by increasing the probability of inhibitory transmitter release.

**Figure 5.**
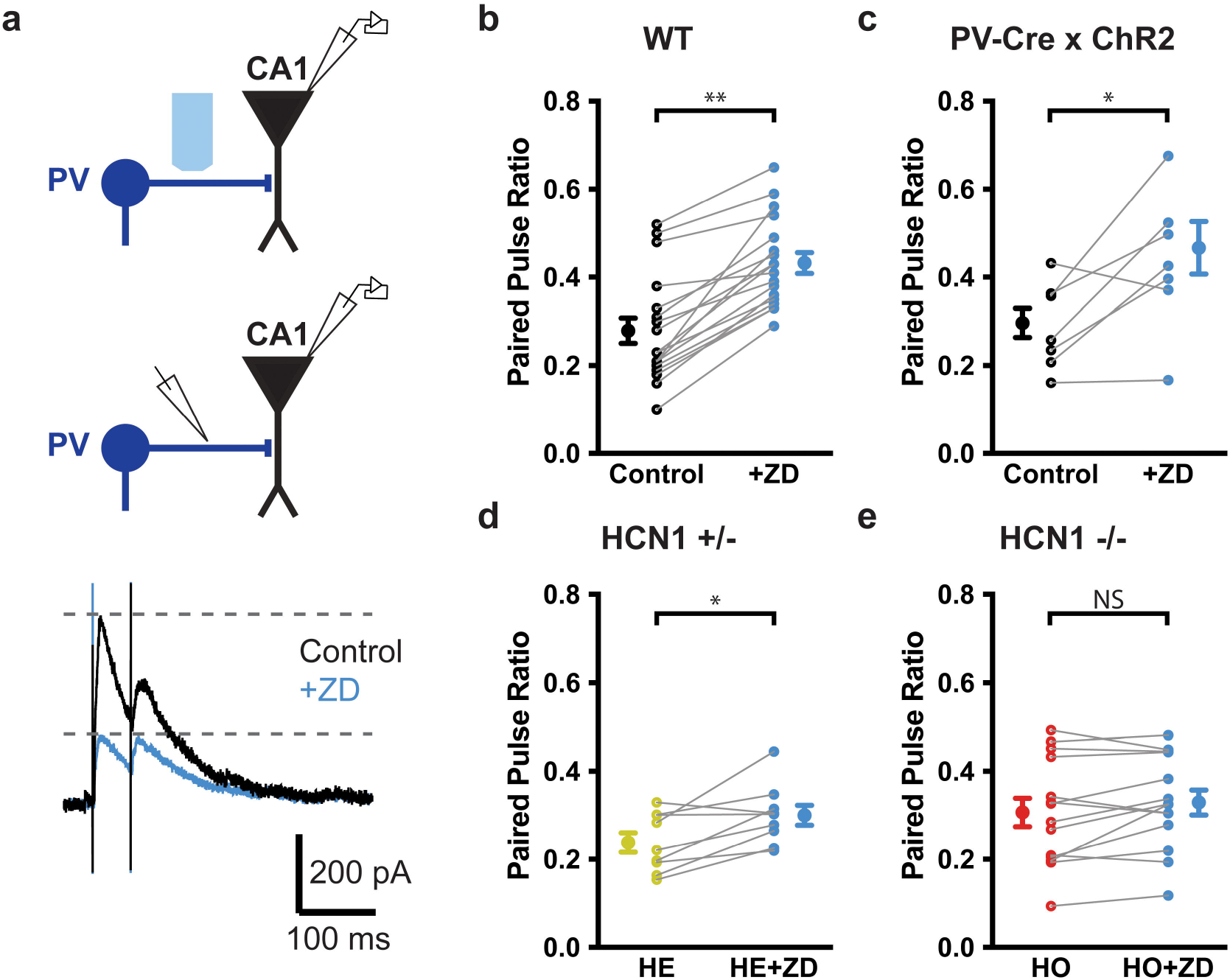
HCN channel blockade increases paired pulse ratio of IPSCs evoked by PV+ IN stimulation. **(a)** (top) Schematic of voltage clamp recordings from CA1 pyramidal neurons with a stimulating electrode in the pyramidal cell layer or optogenetic stimulation using blue (470 nm) light pulses (2 ms). (bottom) Example trace of IPSCs in response electrical 20 Hz paired pulse stimulation before (black) and after (blue) ZD7288 (10 μM) application. Excitatory transmission was blocked with CNQX (25 μM) and APV (50 μM). QX-314 was used in the recording pipette to block postsynaptic HCN channels. **(b)** Paired pulse ratio (PPR) before (black) and after (blue) bath application of ZD7288 in response to 20 Hz extracellular electrical stimulation. **(c)** Paired pulse ratio (PPR) before (black) and after (blue) bath application of ZD7288 in response to 20 Hz light stimulation of ChR2 expressing PV+ IN axon terminals. **(d)** PPR before (yellow) and after (blue) bath application of ZD7288 in response to 20 Hz extracellular electrical stimulation in HCN^+/-^ mice. **(e)** PPR before (red) and after (blue) bath application of ZD7288 in response to 20 Hz extracellular electrical stimulation in HCN^+/+^ mice.

### HCN channels promote activation of individual presynaptic boutons in PV+ INs

Next we turned to calcium imaging to observe more directly the effect of HCN channel blockade on the activation of PV+ IN boutons during electrical stimulation. We injected a Cre-dependent AAV to express an axon-targeting genetically encoded fluorescent calcium indicator (*AAV5-hSynapsin1-FLEx-axon-GCaMP6s*) into area CA1 of PV-Cre mice (Figure 6a). We then used two-photon microscopy to image calcium transients, measured as △F/F, within the PV+ IN axon boutons within the CA1 SP layer in acute brain slices (Figure 6b). We first activated the terminals using three separate trains of stimuli, with each train consisting of 5 pulses applied at 30 Hz, with a 15 s interval between trains. All recordings were performed in the presence of CNQX and APV, first in the absence and then in the presence of ZD7288.

**Figure 6.**
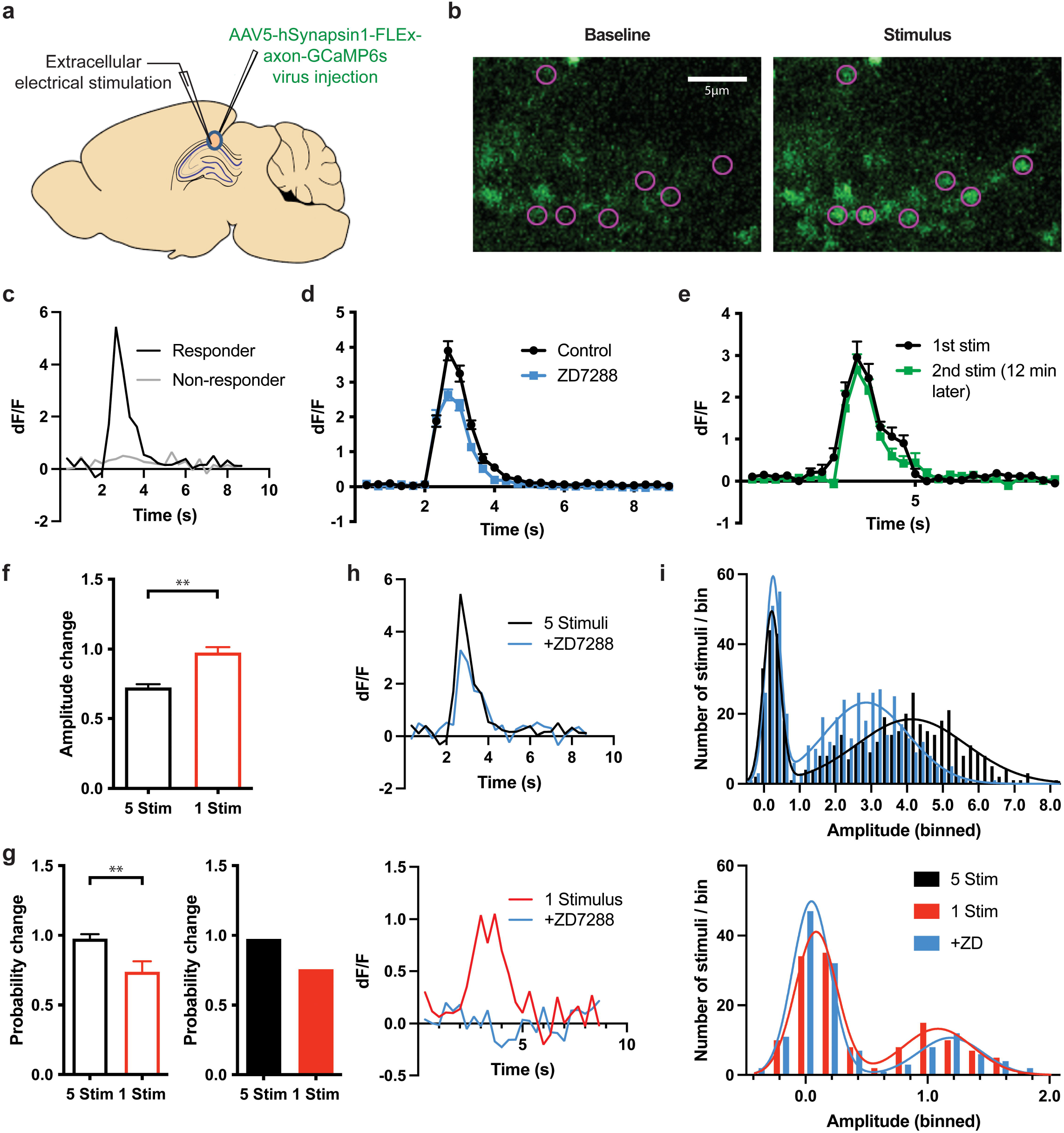
Two-photon imaging of PV+ IN axon-specific GcaMP6s in acute brain slices. **(a)**Schematic representation of viral injection site of axon-GCaMP6s and the location of in vitro extracellular electrical stimulation. **(b)** Sample images of boutons before (left) and during (right) a stimulus response; active boutons are marked by pink circles. **(c)** ΔF/F of two example boutons in response to 5 pulses at 30-Hz extracellular electrical stimulation in the pyramidal cell layer of CA1 at t = 2 s; one bouton was classified as responding (black) and the other as non-responding (grey). **(d)** Average ΔF/F of responding boutons before (black) and after (blue) application of ZD7288. **(e)** Average ΔF/F of responding boutons shortly (< 2 min) after start of imaging (black) and 12 minutes later (green). **(f)** Change in ΔF/F amplitude after bath application of ZD7288 using 5 pulses of 30-Hz stimulation (black) or one stimulus pulse only (red). **(g)** Change in response probability after bath application of ZD7288 using 5 pulses of 30-Hz electrical stimulation (black) or one stimulus pulse only (red). Left (outlined bars): Response probability was determined by fitting the response distribution with two Gaussian components and calculating the fractional area under the component corresponding to successes (non-zero peak component). **(h)** ΔF/F of example boutons in response to 5 pulses of 30-Hz stimulation (top, black) or one stimulus pulse only (bottom, red). Blue traces show the ΔF/F of the same boutons to the same stimulation protocols after bath application of ZD7288. **(i)** Distribution of peak ΔF/F amplitudes in response to either 5 pulses of 30-Hz stimulation (top, black, n = 486 stimuli in 54 boutons) or one stimulus pulse (bottom, red, n = 135 stimuli in 15 boutons) of extracellular electrical stimulation in the pyramidal cell layer. Blue bars show the ΔF/F amplitude distribution of the same boutons to the same stimulation protocols after bath application of ZD7288. Lines show best double gaussian fits with two Gaussian components.

We found that a fraction of the labeled boutons in a given field responded to the electrical stimulation with a transient increase in fluorescence intensity (Figure 6c). The fluorescence response of certain boutons fluctuated from train to train, with some trains eliciting a noticeable △F/F transient and other trains failing to elicit a response. We analyzed the subset of responsive boutons, defined as any bouton in which the △F/F transient elicited by at least one of the three trains of stimuli was greater than 3 times the standard deviation of the baseline fluorescence, with the transient measured within a 1.6 second time window after stimulation. We found that the trains of synaptic stimulation typically activated 20.4 ± 2.6% of all labeled boutons (range 7-42%; n= 1076 boutons from 16 slices from 8 animals). The mean peak △F/F amplitude of the Ca^2+^ transients in these responding boutons was reduced by ZD7288 to 68% of its initial value, from 4.1 ± 0.3 to 2.8 ± 0.2 (p = 0.0004; n = 219 boutons from 16 slices from 8 animals; Figure 6d). This decrease was not due to a rundown of the response during the second train as there was no significant decrease in the size of the calcium transient when we gave two trains of stimuli over an identical time frame in the absence of ZD7288 (△F/F = 2.95 ± 0.38, compared to 2.66 ± 0.37 after 10 minutes; p = 0.36; n = 123 boutons from 11 slices from 4 animals; Figure 6e). These results indicate that HCN1 blockade directly reduces Ca^2+^ influx into the PV+ IN presynaptic boutons.

The effect of ZD7288 could result from a reduction in the size of the Ca^2+^ transient elicited by a single action potential or a decrease in the number of action potentials that invade the presynaptic bouton in response to the brief train of electrical stimulation. In a first attempt to distinguish between these possibilities, we analyzed the individual Ca^2+^ responses of single boutons to each train of stimuli, first in the absence and then in the presence of ZD7288. We characterized the response to each train as a “success” if the response was above the three standard deviation threshold and a “failure” if the response was below the threshold. If HCN channel block decreased the probability that a bouton was activated by an action potential, we predicted that we would observe an increase in the fraction of responses that were “failures”. We found that the vast majority of responses in the absence of ZD7288, 70.8%, were “successes”. Application of ZD7288 decreased the mean ΔF/F peak amplitude of the “successes” in individual boutons to 72.3 ± 2.4% of their amplitude in the absence of the drug (p < 0.0001; n = 486 stimuli in 56 boutons from 4 slices from 4 animals; Figure 6f). In contrast, we observed little change in the fraction of failures in response to ZD7288, with the probability of a failure response in ZD7288 equal to 97.8 ± 6.4% of the initial failure rate (Figure 6g left).

Although the above results suggest that ZD7288 may decrease Ca^2+^ influx in response to a given action potential, our use of trains with five stimuli could mask any increase in failure probability, since a “success” might occur even if the train succeeded in eliciting only one or two action potentials in a bouton. We therefore examined the effect of ZD7288 application on the responses of individual boutons to a single electrical stimulus pulse. We applied 9 single pulses, separated from each other by >15 seconds, before and after ZD7288 application. We found that the amplitude of the Ca^2+^ transients elicited by the single stimulus fluctuated between successes and failures, in a roughly all-or-none manner. Prior to application of ZD7288 the probability of success was only 33.7%, much lower than observed with the trains of 5 stimuli. Application of ZD7288 caused a significant 26.3 ± 7.6% decrease in the probability of a success, to 25.6% (p = 0.0039; n = 135 trials in 15 boutons from 4 animals; figure 6g). In contrast, ZD7288 caused no measurable change in the mean peak amplitude of the successful trials (97.6 ± 3.9%; p = 0.77; Figure 6f). These results indicate that HCN1 channels increased the probability that an action potential invaded the presynaptic bouton, without altering the Ca^2+^ response when an action potential in the presynaptic bouton did occur.

As our use of the three standard deviation threshold to define successes and failures is somewhat arbitrary, we next examined the amplitude histograms for responses of individual boutons, both to the 5-pulse train and to the single pulses of electrical stimulation (see example traces in Figure 6h). In both cases, the distribution of responses had two well defined peaks of ΔF/F values, with one centered around 0, reflecting the failures, and a second broader peak shifted to positive values, corresponding to the successes (Figure 6i). The histograms in response to the trains or single stimuli were well fit by two Gaussian components (R^2^ > 0.91 for all double gaussian fits), allowing us to define the mean ΔF/F values of each component, their standard deviation, and fractional area under the two components, which corresponds to the probability of observing a success or failure. With the 5-pulse train, the fraction of the area under the failure peak was quite low (28.7%). Application of ZD7288 caused a 30.3% decrease in the peak value of the success component (from 4.17 before ZD7288 to 2.86 after ZD7288), without noticeably changing the fraction of successes or failure (71.3% success rate before ZD7288, compared to 69.5% after ZD7288). In contrast, with single stimuli, we observed a smaller fraction of successes (35.3%) compared to the responses to the trains of 5 stimuli. Importantly, application of ZD7288 caused little change in the ΔF/F value at the peak of the successes (ΔF/F = 1.09 before ZD7288, compared to 1.19 after ZD7288), but decreased the fraction of successes by 24.4% (Figure 6i, bottom and 6g, right). These data are in agreement with the results using a simple threshold to classify responses into success and failures, and support the view that blockade of HCN channels reduced the probability that a bouton will respond to an electrical stimulus, without altering the amplitude of the Ca^2+^ response when a bouton did respond to the stimulus.

## Discussion

In this study, we found that HCN1 channels are enriched in the axon terminals of PV+ interneurons in the CA1 region of mouse hippocampus, where they function to enhance inhibitory synaptic transmission by promoting the evoked release of GABA. Prior immunohistochemical labeling experiments, both at the light microscopy and electron microscopy level, have shown that HCN1 subunits are expressed at very high levels in the axonal terminals of PV+ INs throughout the brain, including the hippocampus and neocortex (Bender et al., 2007; Lujan et al., 2005; Notomi and Shigemoto, 2004). However, prior to our study, there has been no direct examination of the role of HCN1 in presynaptic function of PV+ INs.

Here we used a combination of pharmacological, genetic knockout, and optogenetic approaches to examine specifically the role of HCN1 in PV+ INs, in both determining the somatic membrane properties and in modulating evoked inhibitory synaptic transmission onto CA1 PNs. Our whole-cell patch clamp recordings from PV+ IN somata confirmed that HCN channels have a minimal impact on somatic passive and active membrane properties (Figure 2). This is consistent with our triple immunohistochemical labeling, which showed that HCN1 and PV colocalization was restricted to the axon and presynaptic boutons. It also confirms previous studies that reported predominantly axonal and presynaptic expression of HCN channels in hippocampal PV+ INs (Aponte et al., 2006; Elgueta et al., 2015; Roth and Hu, 2020).

Although most PV+ INs did not show a prominent somatic voltage sag in response to hyperpolarizing current steps, a characteristic signature of Ih, a minority of PV+ neurons did show a modest ZD7288-sensitive somatic sag. This may reflect the fact that there are several subclasses of PV+ INs, including soma-targeting basket cells, axon initial segment targeting chandelier cells and dendrite targeting bistratified cells (Booker and Vida, 2018; Klausberger and Somogyi, 2008; Maccaferri, 2005; Que et al., 2021) and these may differ in their expression of HCN channels.

Our studies also provide insight into the mechanism by which HCN1 enhances the inhibitory postsynaptic response. Both through measures of the paired-pulse ratio and Ca^2+^ imaging from PV+ IN presynaptic boutons, our results indicate that presynaptic HCN1 channels function to enhance GABA release. In addition, our imaging experiments further indicate that HCN1 does not enhance the presynaptic Ca^2+^ response to a presynaptic action potential, but rather acts to enhance the probability that an electrical stimulus evokes a presynaptic action potential. We therefore conclude that the activation of HCN1 interacts with the Na^+^-dependent AP as it propagates down the axon and into the presynaptic bouton, thus increasing the efficacy of GABA release.

Our findings are consistent with a number of studies that have found that HCN channels contribute to axonal excitability. In particular HCN channels have been found to be important for maintaining persistent firing in both soma and axons, largely through an action to oppose membrane hyperpolarization in response to a train of spikes (Ballo et al., 2012; Byczkowicz et al., 2019; Elgueta et al., 2015; Roth and Hu, 2020; Soleng et al., 2003). Furthermore, in dentate gyrus PV+ basket cells, blockade of HCN channels increases the threshold for antidromic action potentials evoked by even a single electrical stimulus, which also is consistent with an effect of HCN channels to enhance axonal excitability (Aponte et al., 2006). All of these results are consistent with our findings that HCN1 channels appear to increase the probability that an extracellular electrical stimulus elicits an action potential that propagates to PV+ IN presynaptic boutons to trigger Ca^2+^ influx. However, HCN channels at the axon initial segment have been found to increase the threshold for spike initiation (Ko et al., 2016).

In contrast to the relatively consistent finding that axonal HCN channels enhance spike initiation and propagation, reports on the effect of HCN channels in regulating inhibitory synaptic transmission have been more variable. Thus, HCN channels have been found to enhance the frequency, but not amplitude, of miniature inhibitory postsynaptic currents (mIPSCs) recorded in dentate gyrus granule cells (Aponte et al., 2006). Whereas HCN channels do not affect mIPSC frequency recorded in CA1 pyramidal neurons, they do enhance the frequency of spontaneous, action-potential-dependent IPSCs, with no effect on IPSC amplitude (Lupica et al., 2001). In cerebellum, HCN channels enhance both the frequency and amplitude of spontaneous IPSCs (Southan et al., 2000). In contrast to the above findings that HCN channels promote inhibitory transmitter release, HCN channels decrease mIPSC frequency in globus pallidus (Boyes et al., 2007) and reduce the frequency of both mIPSCs and sIPSCs in medial prefrontal cortex (Cai et al., 2022). An inhibitory effect of HCN channels has also been reported in subsets of excitatory synaptic terminals in entorhinal cortex (Huang et al., 2009; Shah et al., 2004).

Such diverse effects of HCN channels on synaptic transmission could reflect a differential role of the distinct HCN channel subunits expressed in different presynaptic boutons or a differential effect that the same subunit may exert in distinct sets of presynaptic boutons from different populations of inhibitory or excitatory neurons. Some of the conflicting results may also arise from potential off-target effects of ZD7288 on transmitter release (Chevaleyre and Castillo, 2002). To guard against such off-target effects we have limited our application of ZD7288 to a concentration of 10 μM for no more than 10 min. Moreover, by examining mice with a genetic deletion of HCN1, we have been able to verify that the effects we observe with this drug are specific to blockade of HCN1. Finally, by using the PV-Cre mouse line we have further verified that the effects we characterized are mediated by HCN1 and its specific blockade. Future studies in other systems using such genetic-based specific approaches may help resolve some of the discrepant findings.

Our results also have interesting implications for understanding PV+ interneuron function. These inhibitory neurons are known to play an important role in regulating and balancing excitability in the hippocampal network, through feed-forward and feedback inhibition (Geiger et al., 1997; Pouille and Scanziani, 2001; Sohal et al., 2009; Sun et al., 2014). Therefore, the changes in the efficacy of inhibitory GABA release we have observed during pharmacological blockade or genetic deletion of HCN1 are likely to impact network function and animal behavior.

Changes in inhibitory control of network activity are also likely to induce neuropathology. Indeed, impairments in PV+ IN function have been associated with a number of neuronal and cognitive disorders (Jiang et al., 2016; Kann, 2016). For example, alterations in PV+ IN firing due to mutations in the SCNA1 gene that encodes the PV+ IN Nav1.1 voltage-gated Na^+^ channel have been shown to underlie the seizures and intellectual disability seen in individuals with Dravet syndrome (Cheah et al., 2012; Guerrini, 2012; Tai et al., 2014).

Given the importance of PV+ INs for normal brain function, we hypothesize that HCN1 expression in these neurons plays an important role in both the normal function of these neurons in regulating brain activity and in PV+ IN dysfunction in neurological and psychiatric disorders. Recent human genetic studies have identified more than 40 de novo mutations in the HCN1 gene in patients suffering from a range of early childhood onset seizures, ranging from severe early infantile epileptic encephalopathy (EIEE) to generalized epilepsy with febrile seizures (GEFS+) to milder febrile seizure (FS) phenotypes (Marini et al., 2018; Nava et al., 2014; Parrini et al., 2017). Patients carrying the more severe HCN1 mutations also show cognitive impairments and autistic traits. While recent studies have shown the impact of function-impairing HCN channel mutations in pyramidal cells in the cortex and hippocampus (Bleakley et al., 2021; Merseburg et al., 2022) on epileptiform activity, the reduction of inhibition, reported in this study can be expected to contribute to excessive excitability in the hippocampal network and investigating inhibitory inputs from PV+ INs in these pathological mouse lines may be indicated.

## Methods

### Animals

Experiments were conducted on male C57/BL6 mice (Jackson Labs, stock #000664) aged 2-10 months. All animal experiments were conducted in accordance with policies of the NIH Guide for the Care and Use of Laboratory Animals and the Institutional Animal Care and Use Committee (IACUC) of Columbia University. The following commercially available mouse lines were used: *Pvalb^tm1(cre)Arbr^/J* (Jackson Labs, stock #017320); *Gt(ROSA)26Sor^tm14(CAG-tdTomato)Hze^/J* (Jackson Labs, stock #007914); *Gt(ROSA)26Sor^tm32(CAG-COP4*H134R/EYFP)Hze^/J* (Jackson Labs, stock #024109); and *Hcn1^tm2Kndl^/J* (Jackson Labs, stock #016566).

### Immunohistochemistry

Animals were perfused with phosphate buffered saline (PBS) followed by 4% paraformaldehyde in PBS, and brains post-fixed overnight at 4 °C. After several washes in PBS, 40 μm coronal slices were cut using a vibratome, and free-floating sections permeabilized in PBS + 0.1% Triton, followed by incubation in blocking solution (PBS + 5% normal donkey serum) for 1 h at room temperature. Primary antibody incubation was carried out in blocking solution overnight at 4°C. Antibodies used were: mouse monoclonal anti-HCN1 (clone N70-28, NeuroMab 75-110, dilution 1:300; Davis, CA); mouse monoclonal anti-HCN2 (clone N71-37, NeuroMab 75-111, dilution 1:250; Davis, CA); mouse monoclonal anti-Syt2 (Znp-1, Developmental Studies Hybridoma Bank, dilution 1:250; Iowa City, IA); rabbit anti-Parvalbumin (Synaptic Systems 195002, dilution 1:700). Secondary antibody incubation was performed in blocking solution for 2 h at room temperature. All secondary antibodies were used at 1:500 dilutions: goat anti-mouse IgG1 cross-adsorbed (Alexa Fluor 488, Life Technologies A21121; Eugene, OR), goat anti-mouse IgG2a cross-adsorbed (Alexa Fluor 647, Life Technologies, A21241; Eugene, OR), goat anti-rabbit IgG (H+L) cross-adsorbed (Alexa Fluor 568, Life Technologies, A11011; Eugene, OR).Images were acquired on a Zeiss LSM 700 laser scanning confocal microscope with Zen 2012 SP5 FP3 black edition software, using either a Zeiss Fluar 5x/0.25 objective (0.5 zoom, pixel size: 2.5 x 2.5 μm^2^) or a Zeiss Plan-Apochromat 20X/0.8 objective (1.0 zoom, pixel size: 0.3126 x 0.3126 μm^2^).

### Slice preparation

Mice were anesthetized by inhalation of isoflurane (5%) for 7 min, subjected to cardiac perfusion of ice-cold oxygenated artificial cerebrospinal fluid, modified for dissections (d-ACSF; 195 mM sucrose, 10 mM glucose, 10 mM NaCl, 7 mM MgCl_2_, 0.5 mM CaCl_2_, 25 mM NaHCO3, 2.5 mM KCl, 1.25 mM NaH2PO4, 2 mM Na-pyruvate, pH 7.2) for 30 s before decapitation according to the procedures approved by the IACUC of Columbia University. The skull was opened, and the brain removed and immediately transferred into ice-cold carbogenated d-ACSF. The hippocampus was dissected in both hemispheres. Each hippocampus was placed in the groove of an agar block and 400 μm thick hippocampal slices were cut using a vibrating tissue slicer (VT 1200, Leica, Germany) and transferred to a chamber containing a carbogenated mixture of 50% d-ACSF and 50% ACSF (22.5 mM glucose, 125 mM NaCl, 1 mM MgCl2, 2 mM CaCl2, 25 mM NaHCO3, 2.5 mM KCl, 1.25 mM NaH2PO4, 3 mM Na-pyruvate, 1mM ascorbic acid, pH 7.2) at 35°C, where they were incubated for 40 - 60 min. Thereafter, slices were held at room temperature (21°C) until transfer into the recording chamber.

### Slice Electrophysiology

Slices were transferred from the incubation chamber into the recording chamber of an Olympus BX51WI microscope (Olympus, Japan), where they were held in place by a 1mm grid of nylon strings on a platinum frame. Slices were continually perfused with ACSF at 34 ± 1°C, maintained by a thermostat-controlled flow-through heater (Warner Instruments, CT, USA). Healthy somas of CA1 pyramidal neurons were identified visually under 40X (20 x 2) magnification and patched under visual guidance using borosilicate glass pipettes (I.D. 0.75 mm, O.D. 1.5 mm, Sutter Instruments, UK) with a tip resistance of 4 −5.5 MΩ, connected to a Multiclamp 700B amplifier (Molecular Devices, CA) and filled with intracellular solution, containing (in mM): 135 K-gluconate, 5 KCl, 0.1 EGTA, 10 HEPES, 2 NaCl, 5 MgATP, 0.4 Na2GTP, 10 Na2-Phosphocreatin, adjusted to a pH of 7.2 with KOH. Recordings were only accepted if the series resistance after establishing a whole-cell configuration did not exceed 25 MΩ and did not change by more than 20% of the initial value during the course of the experiment.

Extracellular electric stimulation was performed by inserting a borosilicate glass pipette filled with 1M KCl into stratum pyramidale at least 150 μm from the patching site, connected to a custom built variable power supply and triggered via a TTL pulse through the output channels of the recording software (see below). Light stimulation of ChR2 expressing axon terminals was achieved by using a 470 nm pE-100 excitation light source (CoolLED, UK), connected to the fluorescent light path of the microscope, also triggered via TTL pulse.

Pharmacology: Stock solution of 10 mM ZD7288 (Tocris, UK) was stored at −20 °C and diluted in ACSF to a concentration of 10 μM before bath application to the slice. In some experiments, blockers of AMPA and NMDA receptors, 25 μM CNQX and 50 μM APV, respectively, were added to the bath solution and ivabradine was added to the bath solution at a concentration of 30 μM.

### Stereotaxic virus injection

Mice were anaesthetized using isoflurane (Covetrus, Portland, ME) and provided analgesics (Carprofen, Zoetis, Troy Hills, NJ). A craniotomy was performed above the target region and a glass pipette was stereotaxically lowered to the desired depth. Injections were performed using a nano-inject II apparatus (Drummond Scientific), with 25 nl of solution delivered every 15 s until a total amount of 200 nl was reached. The pipette was retracted after 5 min. *AAV-hSynapsin1-FLEx-axon-GCaMP6s* virus (Addgene, MA, USA) was injected bilaterally, at a titer of 1×10^12^ vg/ml, with injection coordinates AP −1.55, ML +/−1.05, DV −1.5 (in millimeters with Bregma as reference). One single virus injection was performed per hemisphere, and each hemisphere counted as an independent injection site.

### 2 photon imaging

Slices were treated in the same way as they were for electrophysiological recordings (see above) and were then transferred into a recording chamber under an Olympus BX61WI microscope at 40x magnification (2 times 20x) connected to a Prairie imaging system, using a Deep See 2-photon laser (Spectra-Physics, CA, USA), exciting GCaMP6s at a frequency of 820 nm. Extracellular electric stimulation was performed by inserting a borosilicate glass pipette filled with 1M KCl into *stratum pyramidale*, connected to a custom-built variable power supply and triggered via a TTL pulse through the output channels of the recording software. Electrical stimulation and 2-photon imaging were also synchronized via a TTL pulse. Slices were scanned manually for boutons, excitable by a 30 Hz 5 pulse extracellular stimulus train. A region of interest was set and plane scans were performed of a small area around the selected boutons, such that the image sequence captured at a minimum of 3 frames per second.

### Data acquisition and analysis

Electrophysiological recordings were digitized, using a Digidata 1322A A/D interface (Molecular Devices, CA), at a sampling rate of 20 kHz (low pass filtered at 10 kHz) and recorded pClamp 10 software (Molecular Devices, CA, USA). The amplifier setting of the Multiclamp 700B were controlled through Multiclamp Commander (Molecular Devices, CA, USA). After 50 ms baseline recording, 1-s current steps of –350 to +350 pA were applied to the patched cells in increments of 25 pA, after which an additional 1s of post-step membrane potential was recorded. The trigger time between these episodes was 3 s. Voltage deflections in response to current steps of –50 to +50 pA were used to calculate the input resistance. Initial resting membrane potential (RMP) was obtained immediately upon breaking into the cell and monitored throughout the recording. Voltage sag in response to negative current steps was calculated by dividing the steady state voltage deflection during the late phase of the –100 pA current step by the peak of the voltage deflection during the same step. Action potential (AP) threshold was determined as the membrane voltage at which the derivative of the voltage trace exceeded 40 mV/ms. Data was analyzed using Axograph X software (Axograph Scientific, Australia), MATLAB (Mathworks, MA, USA) as well as Microsoft Excel (Microsoft Corp., WA, USA) or Prism 8 (Graphpad, CA, USA) and visualized in Acrobat Illustrator (Adobe, CA, USA). To determine statistical significance, paired t-test was used for paired data and unpaired t-test was used for non-piared data, unless otherwise stated.

Confocal microscopy image analysis was performed using ImageJ 1.49v software (National Institutes of Health, USA). 2-photon image sequences were validated by eye, using ImageJ software and further analyzed using custom Matlab scripts. For bouton detection, Images were filtered, using a temporal correlation filter (Pnevmatikakis et al., 2016) and a 2D median filter. Region of interest (ROI) centers for each bouton were determined as local 2D peaks. ROI size was set to a circle around the peak with a radius of 3 pixels (the average size of a bouton at the magnification, used) around the local peak. ΔF/F within these ROIs was then calculated from the original image sequence.

## Acknowledgements

We would like to thank Sami Hassan for help with the 2-photon image analysis and Anastasia Barnett for technical assistance in preparing the immunohistochemical staining. This work was supported by grant R01NS123648 from the NIH (PI, SAS).

## References

Aponte, Y., Lien, C.C., Reisinger, E., and Jonas, P. (2006). Hyperpolarization-activated cation channels in fast-spiking interneurons of rat hippocampus. J Physiol 574, 229–243.

Ballo, A.W., Nadim, F., and Bucher, D. (2012). Dopamine modulation of Ih improves temporal fidelity of spike propagation in an unmyelinated axon. J Neurosci 32, 5106–5119.

Bender, R.A., Kirschstein, T., Kretz, O., Brewster, A.L., Richichi, C., Ruschenschmidt, C., Shigemoto, R., Beck, H., Frotscher, M., and Baram, T.Z. (2007). Localization of HCN1 channels to presynaptic compartments: novel plasticity that may contribute to hippocampal maturation. J Neurosci 27, 4697–4706.

Benes, F.M. (2015). Building models for postmortem abnormalities in hippocampus of schizophrenics. Schizophr Res 167, 73–83.

Bezaire, M.J., and Soltesz, I. (2013). Quantitative assessment of CA1 local circuits: knowledge base for interneuron-pyramidal cell connectivity. Hippocampus 23, 751–785.

Biel, M., Wahl-Schott, C., Michalakis, S., and Zong, X. (2009). Hyperpolarization-activated cation channels: from genes to function. Physiol Rev 89, 847–885.

Bleakley, L.E., McKenzie, C.E., Soh, M.S., Forster, I.C., Pinares-Garcia, P., Sedo, A., Kathirvel, A., Churilov, L., Jancovski, N., Maljevic, S., et al. (2021). Cation leak underlies neuronal excitability in an HCN1 developmental and epileptic encephalopathy. Brain.

Booker, S.A., and Vida, I. (2018). Morphological diversity and connectivity of hippocampal interneurons. Cell Tissue Res 373, 619–641.

Boyes, J., Bolam, J.P., Shigemoto, R., and Stanford, I.M. (2007). Functional presynaptic HCN channels in the rat globus pallidus. Eur J Neurosci 25, 2081–2092.

Byczkowicz, N., Eshra, A., Montanaro, J., Trevisiol, A., Hirrlinger, J., Kole, M.H., Shigemoto, R., and Hallermann, S. (2019). HCN channel-mediated neuromodulation can control action potential velocity and fidelity in central axons. Elife 8.

Cai, W., Liu, S.S., Li, B.M., and Zhang, X.H. (2022). Presynaptic HCN channels constrain GABAergic synaptic transmission in pyramidal cells of the medial prefrontal cortex. Biol Open 11.

Caroni, P. (2015). Inhibitory microcircuit modules in hippocampal learning. Curr Opin Neurobiol 35, 66–73.

Cheah, C.S., Yu, F.H., Westenbroek, R.E., Kalume, F.K., Oakley, J.C., Potter, G.B., Rubenstein, J.L., and Catterall, W.A. (2012). Specific deletion of NaV1.1 sodium channels in inhibitory interneurons causes seizures and premature death in a mouse model of Dravet syndrome. Proc Natl Acad Sci U S A 109, 14646–14651.

Chevaleyre, V., and Castillo, P.E. (2002). Assessing the role of Ih channels in synaptic transmission and mossy fiber LTP. Proc Natl Acad Sci U S A 99, 9538–9543.

Chiu, C.Q., Lur, G., Morse, T.M., Carnevale, N.T., Ellis-Davies, G.C., and Higley, M.J. (2013). Compartmentalization of GABAergic inhibition by dendritic spines. Science 340, 759–762.

Deng, X., Gu, L., Sui, N., Guo, J., and Liang, J. (2019). Parvalbumin interneuron in the ventral hippocampus functions as a discriminator in social memory. Proc Natl Acad Sci U S A 116, 16583–16592.

Elgueta, C., Kohler, J., and Bartos, M. (2015). Persistent discharges in dentate gyrus perisomainhibiting interneurons require hyperpolarization-activated cyclic nucleotide-gated channel activation. J Neurosci 35, 4131–4139.

Freund, T.F. (2003). Interneuron Diversity series: Rhythm and mood in perisomatic inhibition. Trends Neurosci 26, 489–495.

Fuchs, E.C., Zivkovic, A.R., Cunningham, M.O., Middleton, S., Lebeau, F.E., Bannerman, D.M., Rozov, A., Whittington, M.A., Traub, R.D., Rawlins, J.N., et al. (2007). Recruitment of parvalbumin-positive interneurons determines hippocampal function and associated behavior. Neuron 53, 591–604.

Garcia-Junco-Clemente, P., Cantero, G., Gomez-Sanchez, L., Linares-Clemente, P., Martinez-Lopez, J.A., Lujan, R., and Fernandez-Chacon, R. (2010). Cysteine string protein-alpha prevents activity-dependent degeneration in GABAergic synapses. J Neurosci 30, 7377–7391.

Geiger, J.R., Lubke, J., Roth, A., Frotscher, M., and Jonas, P. (1997). Submillisecond AMPA receptor-mediated signaling at a principal neuron-interneuron synapse. Neuron 18, 1009–1023.

Guerrini, R. (2012). Dravet syndrome: the main issues. Eur J Paediatr Neurol 16 Suppl 1, S1–4.

Huang, Z., Walker, M.C., and Shah, M.M. (2009). Loss of dendritic HCN1 subunits enhances cortical excitability and epileptogenesis. J Neurosci 29, 10979–10988.

Hussaini, S.A., Kempadoo, K.A., Thuault, S.J., Siegelbaum, S.A., and Kandel, E.R. (2011). Increased size and stability of CA1 and CA3 place fields in HCN1 knockout mice. Neuron 72, 643–653.

Jiang, X., Lachance, M., and Rossignol, E. (2016). Involvement of cortical fast-spiking parvalbumin-positive basket cells in epilepsy. Prog Brain Res 226, 81–126.

Kaneko, K., Currin, C.B., Goff, K.M., Wengert, E.R., Somarowthu, A., Vogels, T.P., and Goldberg, E.M. (2022). Developmentally regulated impairment of parvalbumin interneuron synaptic transmission in an experimental model of Dravet syndrome. Cell Rep 38, 110580.

Kann, O. (2016). The interneuron energy hypothesis: Implications for brain disease. Neurobiol Dis 90, 75–85.

Katsarou, A.M., Moshe, S.L., and Galanopoulou, A.S. (2017). Interneuronopathies and Their Role in Early Life Epilepsies and Neurodevelopmental Disorders. Epilepsia Open 2, 284–306.

Kessi, M., Peng, J., Duan, H., He, H., Chen, B., Xiong, J., Wang, Y., Yang, L., Wang, G., Kiprotich, K. et al. (2022). The Contribution of HCN Channelopathies in Different Epileptic Syndromes, Mechanisms, Modulators, and Potential Treatment Targets: A Systematic Review. Front Mol Neurosci 15, 807202.

Klausberger, T., and Somogyi, P. (2008). Neuronal diversity and temporal dynamics: the unity of hippocampal circuit operations. Science 321, 53–57.

Ko, K.W., Rasband, M.N., Meseguer, V., Kramer, R.H., and Golding, N.L. (2016). Serotonin modulates spike probability in the axon initial segment through HCN channels. Nat Neurosci 19, 826–834.

Korotkova, T., Fuchs, E.C., Ponomarenko, A., von Engelhardt, J., and Monyer, H. (2010). NMDA receptor ablation on parvalbumin-positive interneurons impairs hippocampal synchrony, spatial representations, and working memory. Neuron 68, 557–569.

Lapray, D., Lasztoczi, B., Lagler, M., Viney, T.J., Katona, L., Valenti, O., Hartwich, K., Borhegyi, Z., Somogyi, P., and Klausberger, T. (2012). Behavior-dependent specialization of identified hippocampal interneurons. Nat Neurosci 15, 1265–1271.

Lorincz, A., Notomi, T., Tamas, G., Shigemoto, R., and Nusser, Z. (2002). Polarized and compartment-dependent distribution of HCN1 in pyramidal cell dendrites. Nat Neurosci 5, 1185–1193.

Lujan, R., Albasanz, J.L., Shigemoto, R., and Juiz, J.M. (2005). Preferential localization of the hyperpolarization-activated cyclic nucleotide-gated cation channel subunit HCN1 in basket cell terminals of the rat cerebellum. Eur J Neurosci 21, 2073–2082.

Lupica, C.R., Bell, J.A., Hoffman, A.F., and Watson, P.L. (2001). Contribution of the hyperpolarization-activated current (I(h)) to membrane potential and GABA release in hippocampal interneurons. J Neurophysiol 86, 261–268.

Maccaferri, G. (2005). Stratum oriens horizontal interneurone diversity and hippocampal network dynamics. J Physiol 562, 73–80.

Magee, J.C. (1998). Dendritic hyperpolarization-activated currents modify the integrative properties of hippocampal CA1 pyramidal neurons. J Neurosci 18, 7613–7624.

Magee, J.C. (1999). Dendritic lh normalizes temporal summation in hippocampal CA1 neurons. Nat Neurosci 2, 508–514.

Marini, C., Porro, A., Rastetter, A., Dalle, C., Rivolta, I., Bauer, D., Oegema, R., Nava, C., Parrini, E., Mei, D., et al. (2018). HCN1 mutation spectrum: from neonatal epileptic encephalopathy to benign generalized epilepsy and beyond. Brain 141, 3160–3178.

Merseburg, A., Kasemir, J., Buss, E.W., Leroy, F., Bock, T., Porro, A., Barnett, A., Troder, S.E., Engeland, B., Stockebrand, M., et al. (2022). Seizures, behavioral deficits, and adverse drug responses in two new genetic mouse models of HCN1 epileptic encephalopathy. Elife 11.

Murray, A.J., Sauer, J.F., Riedel, G., McClure, C., Ansel, L., Cheyne, L., Bartos, M., Wisden, W., and Wulff, P. (2011). Parvalbumin-positive CA1 interneurons are required for spatial working but not for reference memory. Nat Neurosci 14, 297–299.

Nava, C., Dalle, C., Rastetter, A., Striano, P., de Kovel, C.G., Nabbout, R., Cances, C., Ville, D., Brilstra, E.H., Gobbi, G., et al. (2014). De novo mutations in HCN1 cause early infantile epileptic encephalopathy. Nat Genet 46, 640–645.

Nolan, M.F., Malleret, G., Dudman, J.T., Buhl, D.L., Santoro, B., Gibbs, E., Vronskaya, S., Buzsaki, G., Siegelbaum, S.A., Kandel, E.R., et al. (2004). A behavioral role for dendritic integration: HCN1 channels constrain spatial memory and plasticity at inputs to distal dendrites of CA1 pyramidal neurons. Cell 119, 719–732.

Nolan, M.F., Malleret, G., Lee, K.H., Gibbs, E., Dudman, J.T., Santoro, B., Yin, D., Thompson, R.F., Siegelbaum, S.A., Kandel, E.R., et al. (2003). The hyperpolarization-activated HCN1 channel is important for motor learning and neuronal integration by cerebellar Purkinje cells. Cell 115, 551–564.

Notomi, T., and Shigemoto, R. (2004). Immunohistochemical localization of Ih channel subunits, HCN1-4, in the rat brain. J Comp Neurol 471, 241–276.

Parrini, E., Marini, C., Mei, D., Galuppi, A., Cellini, E., Pucatti, D., Chiti, L., Rutigliano, D., Bianchini, C., Virdo, S., et al. (2017). Diagnostic Targeted Resequencing in 349 Patients with Drug-Resistant Pediatric Epilepsies Identifies Causative Mutations in 30 Different Genes. Hum Mutat 38, 216–225.

Pelkey, K.A., Chittajallu, R., Craig, M.T., Tricoire, L., Wester, J.C., and McBain, C.J. (2017). Hippocampal GABAergic Inhibitory Interneurons. Physiol Rev 97, 1619–1747.

Pnevmatikakis, E.A., Soudry, D., Gao, Y., Machado, T.A., Merel, J., Pfau, D., Reardon, T., Mu, Y., Lacefield, C., Yang, W., et al. (2016). Simultaneous Denoising, Deconvolution, and Demixing of Calcium Imaging Data. Neuron 89, 285–299.

Pouille, F., and Scanziani, M. (2001). Enforcement of temporal fidelity in pyramidal cells by somatic feed-forward inhibition. Science 293, 1159–1163.

Que, L., Lukacsovich, D., Luo, W., and Foldy, C. (2021). Transcriptional and morphological profiling of parvalbumin interneuron subpopulations in the mouse hippocampus. Nat Commun 12, 108.

Rinaldi, A., Defterali, C., Mialot, A., Garden, D.L., Beraneck, M., and Nolan, M.F. (2013). HCN1 channels in cerebellar Purkinje cells promote late stages of learning and constrain synaptic inhibition. J Physiol 591, 5691–5709.

Roth, F.C., and Hu, H. (2020). An axon-specific expression of HCN channels catalyzes fast action potential signaling in GABAergic interneurons. Nat Commun 11, 2248.

Santoro, B., Grant, S.G., Bartsch, D., and Kandel, E.R. (1997). Interactive cloning with the SH3 domain of N-src identifies a new brain specific ion channel protein, with homology to eag and cyclic nucleotide-gated channels. Proc Natl Acad Sci U S A 94, 14815–14820.

Santoro, B., and Shah, M.M. (2020). Hyperpolarization-Activated Cyclic Nucleotide-Gated Channels as Drug Targets for Neurological Disorders. Annu Rev Pharmacol Toxicol 60, 109-131.

Sartiani, L., Mannaioni, G., Masi, A., Novella Romanelli, M., and Cerbai, E. (2017). The Hyperpolarization-Activated Cyclic Nucleotide-Gated Channels: from Biophysics to Pharmacology of a Unique Family of Ion Channels. Pharmacol Rev 69, 354–395.

Schulz, J.M., Knoflach, F., Hernandez, M.C., and Bischofberger, J. (2018). Dendrite-targeting interneurons control synaptic NMDA-receptor activation via nonlinear alpha5-GABAA receptors. Nat Commun 9, 3576.

Shah, M.M., Anderson, A.E., Leung, V., Lin, X., and Johnston, D. (2004). Seizure-induced plasticity of h channels in entorhinal cortical layer III pyramidal neurons. Neuron 44, 495–508.

Sohal, V.S., Zhang, F., Yizhar, O., and Deisseroth, K. (2009). Parvalbumin neurons and gamma rhythms enhance cortical circuit performance. Nature 459, 698–702.

Soleng, A.F., Chiu, K., and Raastad, M. (2003). Unmyelinated axons in the rat hippocampus hyperpolarize and activate an H current when spike frequency exceeds 1 Hz. J Physiol 552, 459–470.

Sommeijer, J.P., and Levelt, C.N. (2012). Synaptotagmin-2 is a reliable marker for parvalbumin positive inhibitory boutons in the mouse visual cortex. PLoS One 7, e35323.

Southan, A.P., Morris, N.P., Stephens, G.J., and Robertson, B. (2000). Hyperpolarization-activated currents in presynaptic terminals of mouse cerebellar basket cells. J Physiol 526 Pt 1, 91–97.

Sun, Y., Nguyen, A.Q., Nguyen, J.P., Le, L., Saur, D., Choi, J., Callaway, E.M., and Xu, X. (2014). Cell-type-specific circuit connectivity of hippocampal CA1 revealed through Cre-dependent rabies tracing. Cell Rep 7, 269–280.

Tai, C., Abe, Y., Westenbroek, R.E., Scheuer, T., and Catterall, W.A. (2014). Impaired excitability of somatostatin-and parvalbumin-expressing cortical interneurons in a mouse model of Dravet syndrome. Proc Natl Acad Sci U S A 111, E3139–3148.

Wulff, P., Ponomarenko, A.A., Bartos, M., Korotkova, T.M., Fuchs, E.C., Bahner, F., Both, M., Tort, A.B., Kopell, N.J., Wisden, W., et al. (2009). Hippocampal theta rhythm and its coupling with gamma oscillations require fast inhibition onto parvalbumin-positive interneurons. Proc Natl Acad Sci U S A 106, 3561–3566.

